# Interphase microtubule disassembly is a signaling cue that drives cell rounding at mitotic entry

**DOI:** 10.1101/2021.06.14.448335

**Authors:** Kévin Leguay, Barbara Decelle, Islam E. Elkholi, Michel Bouvier, Jean-François Côté, Sébastien Carréno

## Abstract

Reorganization of the cortical actin cytoskeleton at mitotic entry is essential to increase membrane tension for cell rounding^1,2^. This spherical shape is necessary for the biogenesis and organization of the mitotic spindle^2-6^. Proteins of the Ezrin, Radixin, Moesin (ERM) family play essential roles in mitotic morphogenesis by linking actomyosin forces to the plasma membrane^2,3,7-10^. While ERMs drive metaphase cell rounding, the cell-cycle signals that prompt their conformational activation in mitosis are unknown^11^. We screened a library of small molecules using novel ERM biosensors^12^ and we unexpectedly found that drugs that disassemble microtubules promote ERM activation. Remarkably, cells disassemble their interphase microtubules while entering mitosis^13^. We further discovered that this disassembly of microtubules acts as a cell-cycle signal that directs ERM activation and metaphase cell rounding. We show that GEF-H1, a Rho-GEF inhibited by microtubule binding, acts downstream of microtubule disassembly to activate ERMs via RhoA and its kinase effector SLK. In addition, we demonstrate that GEF-H1 and Ect2, another Rho-GEF responsible for the generation of mitotic actomyosin forces^6,14^, act together to drive metaphase ERM activation and cell rounding. In summary, we report microtubule disassembly as a cell cycle signal that triggers a signaling network ensuring that actomyosin forces are efficiently integrated at the plasma membrane to promote cell rounding at mitotic entry.

## RESULTS AND DISCUSSION

ERMs exist at the plasma membrane under a closed-inactive and an open-active conformation^12^. Under their closed conformation, their N-terminal four-point-one, ezrin, radixin, moesin (FERM) domain binds to their C-terminal actin-binding domain (C-ERMAD) and ERMs cannot link actin filaments (F-actin) to the plasma membrane^15,16^. Kinases provide a mechanism to open ERMs by phosphorylating a conserved Thr residue (T567, T564 and T558 in Ezrin, Radixin and Moesin, respectively) thereby freeing their C-ERMAD, which is then accessible to interact with F-actin. ERMs are phosphorylated and activated at mitotic entry to integrate F-actin forces at the plasma membrane to increase cortical tension and drive cell rounding^2,3,8^. While the Ser/Thr kinases of the SLK family were found to phosphorylate ERMs^2,3,7-9^, we still do not know what cell-cycle signals prompt their activation when cells enter metaphase^11^.

In order to better understand the regulation of ERMs, we recently designed ERM bioluminescence resonance energy transfer (BRET) based biosensors^12^. Briefly, we fused the bioluminescent donor (*Renilla* luciferase, rLucII) to the C-terminus of individual ERMs. These constructs are anchored at the plasma membrane through a myristoylation and polybasic (MyrPB) motif (MyrPB-E,R,M-rLucII, Fig 1A). The rGFP acceptor is also targeted to the plasma membrane through a prenylation CAAX box (rGFP-CAAX, Fig 1A). Since BRET occurs only when the acceptor and donor are in close proximity (<10 nm)^17^, ERM opening and activation was measured by a decrease in BRET signals.

**Figure 1:**
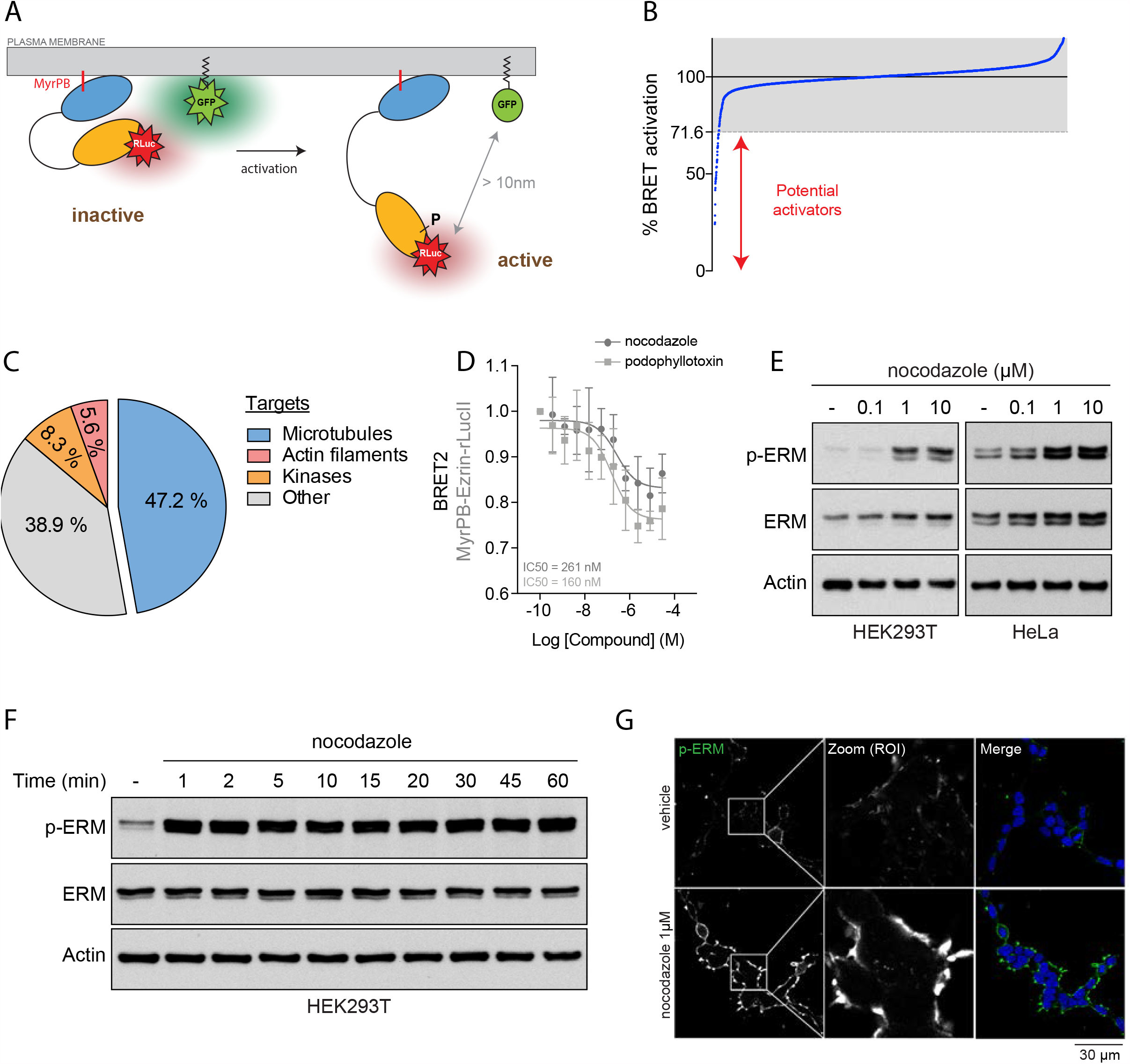
Identification of microtubules as new regulators of ERM proteins. **(A)** Schematic representation of MyrPB-Ezrin-rLucII / rGFP-CAAX bimolecular BRET biosensor^12^. **(B)** Distribution of the 3,469 compounds (blue dots) screened over MyrPB-Ezrin-rLucII BRET activation. BRET2 signals are normalized to vehicle (DMSO, 100%). 71,6% is the threshold of activation that corresponds to 3 times the standard deviation of all compounds. Compounds promoting a BRET2 signal decrease are potential Ezrin activators. **(C)** Diagram representing the identified targets of the validated ERM activators. **(D)** BRET2 signals measured in HEK293T cells expressing the MyrPB-Ezrin biosensor and incubated with increasing concentration of nocodazole (dark grey) or podophyllotoxin (light grey). **(E)** Immunoblot of HEK293T (left) and HeLa (right) cells incubated with the indicated increasing concentration of nocodazole for 15 min. **(F)** Immunoblot of HEK293T cells incubated with 1 µM nocodazole for the indicated times. **(G)** Immunofluorescence of HEK293T cells incubated with vehicle (DMSO, top) or 1 µM nocodazole (bottom) for 15 min. p-ERM (white) and DAPI (blue). BRET2 signals (D) represent the mean +/-s.d. of three independent experiments. Immunoblots (E-F) and immunofluorescences (G) are representative of three independent experiments.

Using the MyrPB-Ezrin-rLucII BRET biosensor, we screened a library of 3,469 FDA-approved small molecules in human embryonic kidney cells (HEK293T). Among the 46 candidates promoting BRET signal decrease (Fig 1B), we identified 36 molecules that activate Ezrin in a confirmation screen (Fig S1A,B). Surprisingly, molecules that promote microtubule disassembly were over-represented among these activators (17 out of 36 molecules; Fig 1C and S1B). By performing dose-response experiments with two different microtubule destabilizing drugs, we found that nocodazole and podophyllotoxin promote Ezrin opening at IC50 compatible with their action on microtubule disassembly (261 nM and 160 nM, respectively, Fig 1D). Next, we confirmed that microtubule disassembly activates ERMs by measuring the phosphorylation of their common regulatory Thr using a well characterized anti p-ERM antibody^3^. We found that microtubule disassembly promotes ERM activation in HEK293T cells as well as in the five other human and murine cell lines tested (Fig 1E and S1C-E). We also found that ERMs are activated after 1 min of nocodazole treatment (Fig 1F) and are phosphorylated at the cell membrane upon microtubule disassembly (Fig 1G and S1F). To our knowledge, this is the first evidence that the membrane-actin linkers ERMs are activated by microtubule disassembly.

We next aimed to determine how microtubule disassembly promotes ERM activation. We previously reported that ERM directly binds to microtubules through a conserved charged motif within their FERM domain^10^. However, we showed that ERM activation by nocodazole is not dependent on their direct interaction with microtubules since an Ezrin microtubule binding mutant (Ezrin^KK211,212MM^) is still phosphorylated after nocodazole treatment (Fig S2A). We then aimed to identify which kinase phosphorylates ERMs upon microtubule disassembly. We first knocked down SLK using two independent shRNA and found that its depletion totally abrogates ERM phosphorylation upon nocodazole treatment (Fig 2A-B). Then, using an *in vitro* kinase assay with immuno-precipitated SLK from cells treated with nocodazole, we showed that microtubule disassembly activates the kinase activity of SLK (Fig 2C-D). Importantly, this was not a direct effect of nocodazole on the enzyme since this molecule did not affect the kinase activity of a recombinant SLK *in vitro* (Fig S2B).

**Figure 2:**
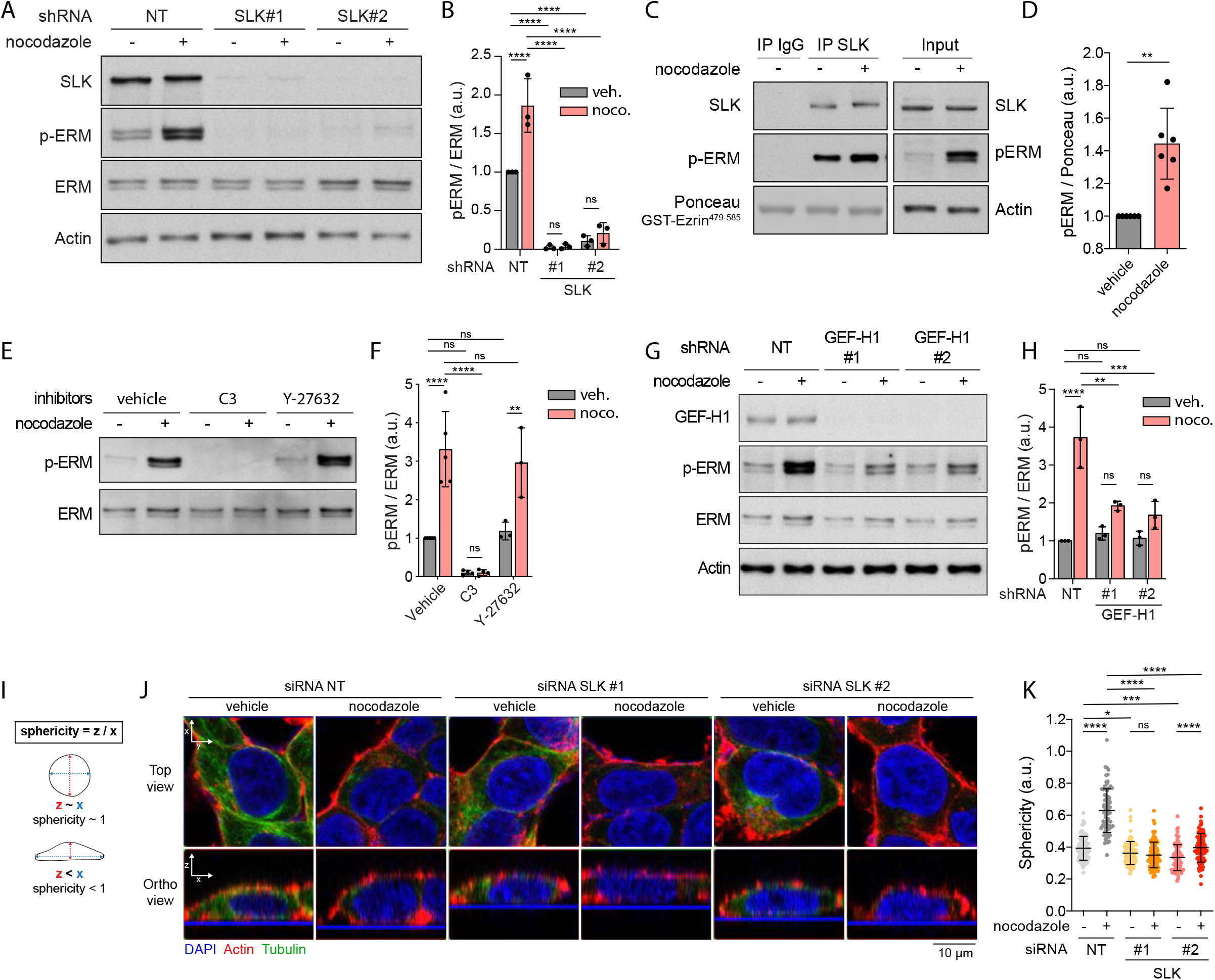
Microtubule disassembly activates ERM proteins through GEF-H1, RhoA and SLK. **(A-B)** Immunoblot of HEK293T cells treated with non-target shRNA (NT) or two independent shRNA targeting SLK and incubated with either vehicle (DMSO) or 1 µM nocodazole for 15 min (A). p-ERM over ERM signals were quantified and normalized to NT incubated with vehicle (B). **(C-D)** Kinase assay of endogenous SLK immunoprecipitated from HEK293T cells incubated with vehicle (DMSO) or 1 µM nocodazole for 15 min. Total lysate (input) is shown on the right (C). p-ERM over GST-Ezrin^479-585^ (substrate) signals measured by Ponceau staining were quantified and normalized to the vehicle (D). **(E-F)** Immunoblot of HEK293T cells pre-incubated with either vehicle (DMSO), 1 µg/mL C3 transferase for 6h or 10 µM Y-27632 for 4h and then incubated with vehicle (DMSO) or 1 µM nocodazole for 15 min (E). p-ERM over ERM signals were quantified and normalized to the vehicle (F). **(G-H)** Immunoblot of HEK293T cells treated with non-target shRNA (NT) or two different shRNA targeting GEF-H1 incubated with either vehicle (DMSO) or 1 µM nocodazole for 15 min (G). p-ERM over ERM signals were quantified and normalized to NT treated with vehicle (H). **(I-K)** Measurement of cell sphericity in HEK293T cells incubated with vehicle (DMSO) or 1 µM nocodazole for 15 min. Cell sphericity (I) was quantified (K) as the ratio between cell height (z) and cell width (x) based on immunofluorescence orthogonal reconstruction of Z stacks (J) (F-actin, red; Tubulin, green; DAPI, blue). Immunoblots (A, C, E, G) and immunofluorescences (J) are representative of at least three independent experiments. P-ERM and sphericity quantifications represent the mean +/-s.d. of at least three independent experiments. Dots represent independent experiments (B, D, F, H) or individual cells (K). P values were calculated using Holm-Sidak’s multiple comparisons test with a single pooled variance (B, F, H, K) or using two-tailed paired t test (D). *, P < 0.05. **, P < 0.01. ***, P < 0.001. ****, P < 0.0001. ns, not significant.

Interestingly, we recently discovered that SLK is a direct effector of RhoA^18^. Here, we found that chemical inhibition of RhoA, using the exoenzyme C3 transferase, abrogates ERM activation after microtubule disassembly (Fig 2E-F). We also found that ROCK, the other kinase effector of RhoA that was shown to phosphorylate ERMs^19^, is not required to activate ERMs downstream of microtubule disassembly since its inhibition using Y-27632 did not decrease ERM phosphorylation after nocodazole treatment (Fig 2E-F and S2C-D).

Nocodazole treatment has been reported to activate RhoA through the release of GEF-H1 from microtubules^20,21^. We established that activation of ERMs following microtubule disassembly was mediated by this Rho-GEF, which is inhibited by its binding to microtubules^21-25^. Depletion of GEF-H1 blocked ERM phosphorylation after nocodazole treatment (Fig 2G-H and S2E-F). Importantly, we noticed that GEF-H1 depletion does not affect ERM phosphorylation in cells with intact microtubules, unlike SLK depletion or RhoA inhibition (Fig 2A-H and S2E-F). This indicates that GEF-H1 activates ERMs only downstream of microtubule disassembly whereas RhoA and SLK also control the global level of ERM activation in resting cells.

Nocodazole treatment was previously found to promote cell rounding in interphase dependently on GEF-H1^20^. We extended these findings by showing that the effect of nocodazole on cell shape is strictly dependent on SLK. Indeed, depletion of SLK prevented the increase of cell sphericity upon microtubule disassembly (Fig 2I-K). Altogether these experiments indicate that microtubule disassembly promotes cell rounding in interphase by activating ERMs through activity of GEF-H1, RhoA and SLK.

Interestingly, at mitotic entry, CDK1 was shown to promote interphase microtubule array disassembly^13^. This frees tubulin dimers that can then re-polymerize to form the mitotic spindle. Treatment with low doses of taxol (paclitaxel), a molecule that stabilizes microtubules, inhibits disassembly of the interphase microtubule array while cells enter mitosis^13^. We treated cells with increasing low dose of taxol and showed that inhibition of interphase microtubule disassembly prevents cell rounding in metaphase as measured by microscopy (Fig 3A-B). In addition, while ERM phosphorylation increases more than threefold in control metaphase cells when compared to interphase, we found that low dose of taxol partially inhibits ERM mitotic activation (Fig 3C-D). Importantly, taxol-induced stabilization of microtubules did not affect the levels of phosphorylated ERMs in interphase (Fig 3D). This suggests that microtubule disassembly at the transition between interphase and mitosis acts as cell-cycle cue that controls ERM activation and mitotic cell rounding.

**Figure 3:**
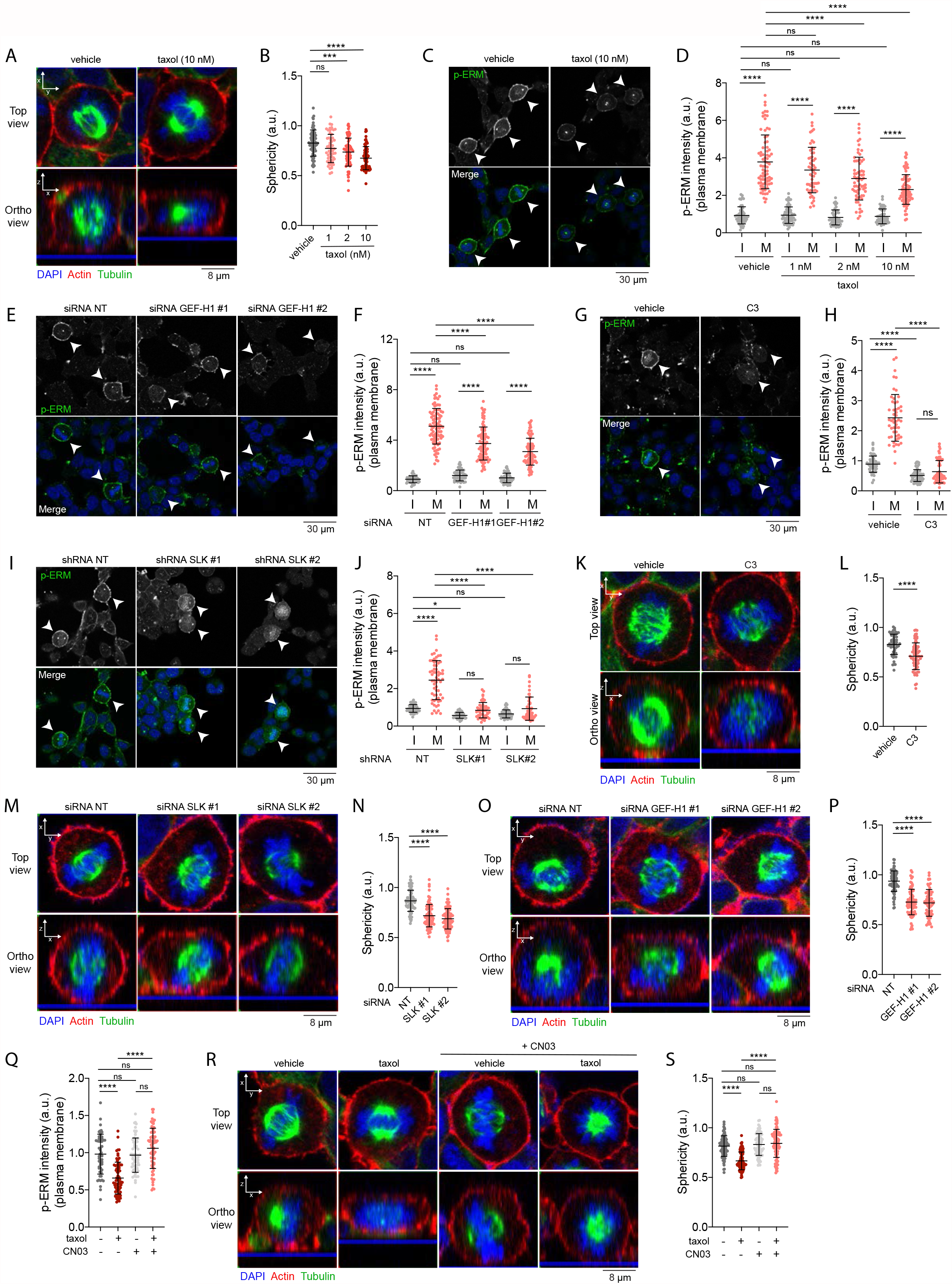
Microtubule disassembly controls ERM activation and cell rounding during mitosis. **(A-B)** Sphericity of metaphase cells (B) was measured on immunofluorescence orthogonal reconstruction of Z stacks (A bottom) (F-actin, red; Tubulin, green; DAPI, blue) in HEK293T metaphase cells incubated with vehicle (DMSO) or increasing concentrations of taxol for 90 min. **(C-J)** p-ERM (white) and DAPI (blue) staining of HEK293T cells incubated with vehicle (DMSO) or increasing concentrations of taxol for 90 min (C-D), treated with non-target siRNA or 2 independent siRNA targeting GEF-H1 (E-F), incubated with vehicle (water) or 1 µg/mL C3 transferase (C3) for 6h (G-H), or treated with non-target shRNA or 2 independent shRNA targeting SLK (I-J). Metaphase and interphase cells were identified based on DAPI staining (C, E, G, I) and p-ERM signal intensity at the plasma membrane is normalized to interphase cells treated with vehicle (D, H) or to interphase cells treated with non-target siRNA/shRNA (NT). I = interphase, M = metaphase. **(K-P)** Sphericity of metaphase cells (L,N,P) was measured on immunofluorescence orthogonal reconstruction of Z stacks (K,M,O bottom) (F-actin, red; Tubulin, green; DAPI, blue) in HEK293T metaphase cells incubated with vehicle (water) or 1 µg/mL C3 transferase (C3) for 6h (K-L), treated with non-target siRNA of 2 independent siRNA targeting SLK (M-N) or treated with non-target siRNA of 2 independent siRNA targeting GEF-H1 (O-P). **(Q)** Quantification of p-ERM staining at the plasma membrane of HEK293T metaphase cells incubated with vehicle (DMSO), 10nM taxol for 90 min and / or 1 µg/mL Rho activator II (CN03) for 1h. Metaphase cells were identified based on DAPI staining (see Fig S3G). P-ERM signal intensity is normalized to cells treated with vehicle. **(R-S)** Sphericity of metaphase cells (S) was measured on immunofluorescence orthogonal reconstruction of Z stacks (R bottom) (F-actin, red; Tubulin, green; DAPI, blue) in HEK293T metaphase cells incubated with vehicle (DMSO), 10nM taxol for 90 min and / or 1 µg/mL Rho activator II (CN03) for 1h. Immunofluorescences are representative of three independent experiments. P-ERM and sphericity quantifications represent the mean +/-s.d. of three independent experiments. Dots represent individual cells (B, D, F, H, J, L, N, P, Q, S). P values were calculated using Sidak’s multiple comparisons test with a single pooled variance. *, P < 0.05. ***, P < 0.001. ****, P < 0.0001. ns, not significant.

We then tested if mitotic disassembly of interphase microtubules could engage the same signaling network that activates ERM phosphorylation and cell rounding after nocodazole-induced microtubule disassembly (see Fig 2). We first established that GEF-H1 is necessary to relay the signal from disassembling microtubules to metaphase ERM activation. Cells depleted for GEF-H1 show a twofold decrease of ERM phosphorylation in metaphase (Fig 3E-F and S3A-B). We also showed that RhoA inhibition and SLK depletion almost totally abrogate phosphorylation of ERMs in metaphase (Fig 3G-J and S3C-D). Interestingly, unlike GEF-H1 depletion, both RhoA inhibition and SLK depletion abrogate ERM activation also in interphase (Fig 3E-J and S3C-D). To our knowledge, GEF-H1 is the first identified protein that promotes ERM activation specifically in mitosis. We then established that GEF-H1 acts with RhoA and SLK to promote rounding of cells entering mitosis. We first confirmed that RhoA is necessary for metaphase cell rounding^14^ by chemically inhibiting the small GTPase using the exoenzyme C3 transferase (Fig 3K-L). We then found that like its Drosophila orthologue Slik^2^, SLK is necessary to promote this shape transformation in human cells (Fig 3M-N). Finally, we discovered that in addition to activating ERMs in metaphase, GEF-H1 is also driving mitotic cell rounding (Fig 3O-P and S3E-F). While GEF-H1 was previously shown to participate in cytokinesis in telophase^26^, this is the first evidence that this Rho-GEF plays roles in the earlier stage of mitosis. If the GEF-H1-dependent pathway transmits signals from interphase microtubule disassembly to ERM activation and cell rounding, we reasoned that requirement of microtubule disassembly at mitotic entry can be bypassed by chemically activating RhoA. We confirmed this hypothesis and found that CN03, a small molecule that activates RhoA, restores normal metaphase levels of p-ERMs and cell sphericity in cells treated by low levels of taxol (Fig 3Q-S and S3G). This demonstrates that disassembly of interphase microtubules acts as cell-cycle cues that engage GEF-H1, RhoA and SLK to promote ERM activation and drive cell rounding in metaphase.

RhoA was shown to be central to another signaling pathway that regulates reorganization of the actin cytoskeleton in early mitosis^6^. RhoA is also activated by Ect2, a Rho-GEF that is itself phosphorylated and activated by Cdk1/Cyclin B, the central complex that drives cells into mitosis^27^. Ect2 was shown to promote the generation of actomyosin forces dependently on the RhoA kinase effector ROCK, which phosphorylates myosin II^6^. Interestingly, while SLK depletion and RhoA inhibition totally abrogated ERM activation in metaphase, GEF-H1 depletion or treatment with low dose of taxol only partially inhibited ERM activation (Fig 3C-J). We therefore hypothesized that interphase microtubule disassembly and Ect2 could act together to fully activate ERMs in metaphase. We first depleted Ect2 and found that this depletion partially decreases ERM phosphorylation in metaphase but did not affect ERM activation in interphase (Fig 4A-B and S4A). This shows that similarly to GEF-H1, Ect2 contributes to ERM activation only in cells entering mitosis. Having confirmed that Ect2 is important for mitotic cell rounding^6^ (Fig. 4C and S4B), we then depleted Ect2 together with GEF-H1 and showed that this co-depletion further decreases ERM phosphorylation in metaphase when compared to single depletions (Fig 4D-E and S4A). We also confirmed this result by depleting Ect2 and inhibiting the GEF-H1 signaling pathway with low dose of taxol to inhibit interphase microtubule disassembly (Fig 4D-E). Consistent with this discovery, we found that these two pathways act together to promote metaphase cell rounding (Fig 4F-G). Ect2 and GEF-H1 depletion led to a similar decrease of cell sphericity in metaphase while their co-depletion further decreased this parameter (Fig 4F-G and S4A). We confirmed this by stabilizing microtubules with low dose of taxol in Ect2 depleted cells (Fig 4F-G). Altogether, this provides evidence that Ect2 and GEF-H1 converge to RhoA to activate ERMs and cell rounding at mitotic entry. Finally, since RhoA directly activates both SLK and ROCK^18^, we aimed to investigate the respective roles of these two kinases in rounding of metaphase cells. As previously reported^14^, we found that ROCK inhibition by Y-27632 decreased cell rounding in metaphase (Fig 4H-I). Yet, we found that SLK depletion promoted a higher decreased sphericity of metaphase cells (Fig 4H-I). Importantly this that was not further potentiated when ROCK was co-inhibited with SLK (Fig 4H-I). This indicates that coupling actin forces to the plasma membrane through SLK-ERM activation plays major roles in metaphase cell rounding.

**Figure 4:**
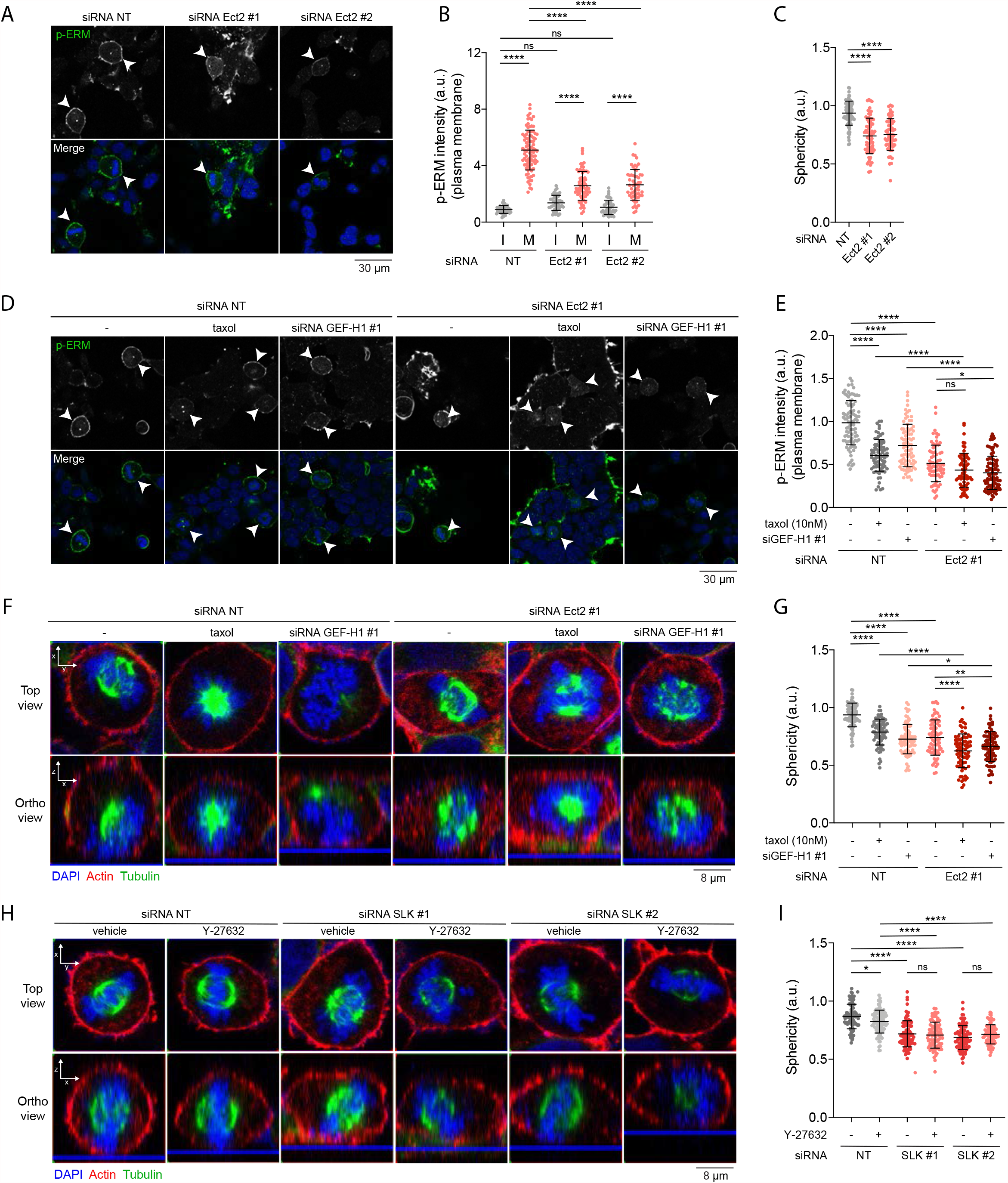
Ect2 and GEF-H1 act together to activate ERM proteins and control cell rounding during mitosis. **(A-B)** p-ERM (white) and DAPI (blue) staining of HEK293T cells treated with non-target siRNA or 2 independent siRNA targeting Ect2 (B). Metaphase and interphase cells were identified based on DAPI staining and p-ERM signal intensity at the plasma membrane is normalized to interphase cells treated with non-target siRNA (NT). I = interphase, M = metaphase. **(C)** Sphericity of metaphase cells was measured on immunofluorescence orthogonal reconstruction of Z stacks (see Fig S4B) in HEK293T metaphase cells treated with non-target siRNA or 2 independent siRNA targeting Ect2. **(D-E)** p-ERM (white) and DAPI (blue) staining of HEK293T cells treated as indicated (D). Metaphase cells were identified based on DAPI staining and p-ERM signal intensity at the plasma membrane is normalized to metaphase cells treated with non-target siRNA (NT). I = interphase, M = metaphase. p-ERM signal intensity of metaphase cells treated with non-target siRNA (NT) or siRNA #1 targeting Ect2 (Ect2 #1) was already shown in Fig 4B. **(F-I)** Sphericity of metaphase cells (G,I) was measured on immunofluorescence orthogonal reconstruction of Z stacks (F,H bottom) (F-actin, red; Tubulin, green; DAPI, blue) in HEK293T metaphase cells treated as indicated (F-H). Sphericity of metaphase cells treated with non-target siRNA (NT) or siRNA #1 targeting Ect2 (Ect2 #1) (G), and metaphase cells treated with non-target siRNA (NT) or siRNA targeting SLK and treated with vehicle (I) was already shown in Fig 4C and 3M-N, respectively. Immunofluorescences (A, D, F, H) are representative of three independent experiments. P-ERM and sphericity quantifications represent the mean +/-s.d. of three independent experiments. Dots represent individual cells (B, C, E, G, I). P values were calculated using Sidak’s multiple comparisons test with a single pooled variance. *, P < 0.05. **, P < 0.01. ****, P < 0.0001. ns, not significant.

Our work identified interphase microtubule disassembly as a cell-cycle cue that controls metaphase cell rounding. This disassembly activates ERMs through GEF-H1, RhoA and SLK (Fig 5). Importantly, interphase microtubule disassembly depends on the inactivation by phosphorylation of Ensconsin/MAP7, a microtubule-stabilizing protein, downstream of Cdk1^13^. Thus, at mitosis onset, the Cdk1/Cyclin B complex directs both Ect2 activation^6^ and interphase microtubule disassembly^13^. Ect2 and GEF-H1 both engage RhoA that relays the cell-cycle signals to its two effectors, ROCK and SLK. ROCK takes in charge the generation of cortical acto-myosin forces^6,14^ while SLK activates ERMs that couple these forces to the plasma membrane (Fig 5). This integrated signaling network ensures that the multiple events that promote the reorganization of the actin cortex are properly coordinated and synergized at mitosis onset to drive cell rounding.

**Figure 5:**
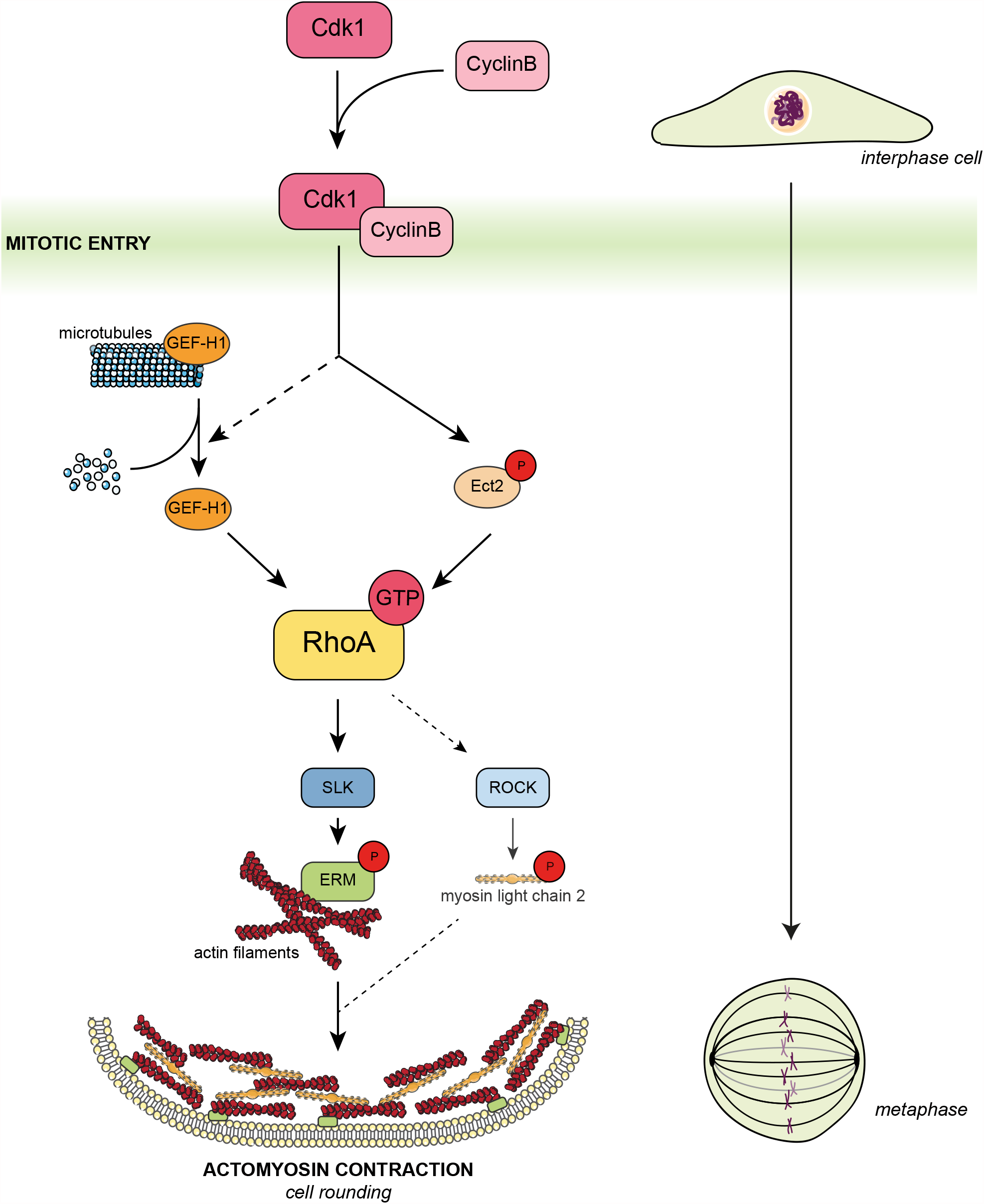
Model for ERM activation and cell rounding at mitotic entry. The Cdk1/cyclin B complex promotes both Ect2 phosphorylation^6^ and interphase microtubule disassembly^13^ leading to GEF-H1 release and activation. Both Rho-GEFs (GEF-H1 and Ect2) activate RhoA that in turn activates SLK and ROCK. While ROCK takes in charge the phosphorylation of myosin II to generate actomyosin forces, SLK activates ERMs that integrate the actomyosin forces to the plasma membrane. This integrated signaling network promotes cell rounding at mitotic entry.

## ACKNOWLEDGMENT

This work has been supported by CCSRI Innovation Grant (705892) to S.C, a project Grant from the CIHR (175193) to S.C. and M.B., a Foundation Grant from the CIHR (148431) to M.B and a a National Science and Engineering Research Council of Canada (RGPIN-201604808) to J.-F.C.

I.E.E. is a recipient of a FRQS Doctoral studentship. K.L held a doctoral scholarship from IRIC and from Montreal University’s Molecular Biology Program as well as a ESP fellowship from Montreal University. MB holds the Canada Research Chair in Signal Transduction and Molecular Pharmacology. J.-F.C. holds the philanthropic Transat Chair in Breast Cancer Research

## MATERIAL & METHODS

### Reagents and inhibitors

Nocodazole, taxol and ROCK inhibitor Y-27632 were purchased from Sigma (#M1404, #T7402 and #688000, respectively). Podophyllotoxin was purchased from Cayman chemical (#19575). The exoenzyme C3 transferase (Rho inhibitor I) and the Rho activator II (CN03) were purchased from Cytoskeleton (#CT04 and #CN03, respectively). Coelenterazine 400a was purchased from NanoLight Technology (#340).

### DNA and RNA constructs

MyrPB-Ezrin-rLucII, Ezrin-rLucII and rGFP-CAAX constructs were previously described^12^. Ezrin^KK211,212MM^-rLucII was obtained by inverse PCR from Ezrin-rLucII using the following primers: forward) 5’-ATGATGGGAACAGACCTTTGGCTTGGAGTTGATGCCCTTGG-3’ and (reverse) 5’-GTTTTTTATCTCGAAATAGTTGATTCCATACATTTCCAGGTCC-3’. MISSION shRNA constructs were obtained in pLKO.1-puro vectors from Sigma: SLK#1 (TRCN0000000895), SLK#2 (TRCN0000000897), GEF-H1#1 (TRCN0000003174), GEF-H1#2 (TRCN0000003176). FlexiTube siRNA were obtained from Qiagen: SLK#1 (SI00107723), SLK#2 (SI04438350), GEF-H1#1 (SI00067564), GEF-H1#2 (SI05181533), Ect2#1 (SI02643067), Ect2#2 (SI03049249).

### Cell culture, transfection and infection

Human HEK293T, HeLa, A375, SW620 cells and murine MC38 cells were cultured in Dulbecco’s modified Eagle’s medium (DMEM; 4.5g/L D-Glucose, L-Glutamine, 110mg/L sodium pyruvate; Invitrogen #11995073) supplemented with 10% fetal bovine serum (FBS; Life Invitrogen #12483020) and 1% penicillin-streptomycin antibiotics (ThermoFisher #15070063) at 37°C with 5% CO_2_. Transfections for BRET experiments were performed as previously described^12^. Briefly, HEK293T cells were transfected with 1µg of total DNA (50ng rLucII construct, 300ng rGFP-CAAX and 650ng salmon sperm DNA) using linear polyethyleneimine (PEI, Alfa Aesar #43896) as transfecting agent (PEI:DNA ratio of 3:1). Transfected cells were then plated in white 96-well culture plates (VWR #82050-736) and incubated for 48h prior BRET measurement. For loss-of-function experiments, siRNAs were transfected using the lipofectamine RNAiMAX (Thermofisher #13778075) as prescribed by the manufacturer. For lentiviral infection, lentiviruses were added to the cells cultivated in DMEM supplemented with 10% FBS and 5µg/ml polybrene (Sigma #H9268) for 48h. Infected cells were then selected for another 48h prior experiments using 2µg/ml puromycin (EMD Millipore #540222).

### BRET measurement

Forty-eight hours after transfection, HEK293T cells were washed with Hank’s balanced salt solution (HBSS, ThermoFisher #14065056) and incubated for 5 min with 2,5µM coelenterazine 400a diluted in HBSS. BRET signals were monitored with a Tecan Infinite 200 PRO multifunctional microplate reader (Tecan) equipped with BLUE1 (370-480nm; donor) and GREEN1 (520-570nm; acceptor) filters. BRET signals were calculated as a ratio by dividing the acceptor emission over the donor emission.

### Chemical screen

The day before the experiment, HEK293T cells stably expressing MyrPB-Ezrin-rLucII and rGFP-CAAX were resuspended in DMEM/F12 without phenol red (Invitrogen #21041025) supplemented with 1% FBS, sifted through a 70 µm-cell strainer (Falcon #352350) and seeded in white 384-well culture plates (Corning #3570). After an overnight incubation at 37°C with 5% CO_2_, compounds from different libraries (Microsource Discovery, Biomol GmbH, Prestwick and Sigma) were added using an Echo 555 acoustic dispenser (Labcyte) at a final concentration of 10 µM for 1h. Substrate solution (coelenterazine 400a diluted in HBSS with 1% pluronic acid) was added 15min before reading at a final concentration of 2.5 µM. BRET signals were monitored with a Synergy NEO HTS microplate reader (BioTek) equipped with a donor filter of 410/80 nm and an acceptor filter of 515/40 nm.

### Immunoblotting

Cells were incubated with the indicated inhibitors for 15min prior lysis, unless mentioned otherwise. After treatment, cells were washed with ice-cold phosphate-buffered saline (PBS) and lysed in TLB buffer (40 mM HEPES, 1 mM EDTA, 120 mM NaCl, 10 mM NaPPi, 10% glycerol, 1% TritonX-100, 0.1% SDS) supplemented with both phosphatase and protease inhibitors (phosphatase inhibitor cocktail (PIC, Sigma #P2850), 1 mM sodium orthovanadate (Na_3_VO_4_, Sigma #S6508), 5 mM β-Glycerophosphate (Sigma #G6251), 1 mM phenylmethylsulfonyl fluoride (PMSF, Sigma #P7626) and anti-protease cocktail (Sigma #4693132001)). Samples were then boiled in sample buffer (200mM TrisHCl 1M pH6.8, 8% SDS, 0.4% bromophenol blue, 40% glycerol and 412mM β-mercaptoethanol) before being resolved by 8% SDS-PAGE and transferred to nitrocellulose membranes (pore 0.2μm, VWR #27376-991). Membranes were blocked in TBS-Tween (25mM Tris-HCl pH 8, 125mM NaCl, 0.1% Tween 20) supplemented with 2% Bovine Serum Albumin (BSA) (Bioshop #ALB001.250) for one hour before overnight incubation with primary antibodies at 4°C. Primary antibodies used are: rabbit anti-ERM (1:1000, Cell Signaling #3142), rabbit anti-Ezrin (1:1000, Cell Signaling #3145), rabbit anti-phospho-ERM (1:5000)^3^, mouse anti-Actin (1:5000, Sigma #MAB1501), rabbit anti-SLK (1:500, Cederlane #A300-499A), rabbit anti-GEF-H1 (1:1000, abcam #ab155785), rabbit anti-Ect2 (1:1000, Millipore Sigma #07-1364) and mouse anti-tubulin (1:1000, Sigma #T9026). Washed membranes were then incubated for 1h with secondary antibodies: goat anti-rabbit HRP antibody (1:10000,Santacruz #sc-2004) or goat anti-mouse HRP (1:10000, Santacruz #sc-516102). Protein detection was performed using Amersham ECL Western Blotting detection reagent (GE Healthcare #CA95038-564L). Immunoblots were quantified using ImageJ software (NIH).

### Immunofluorescence

Cells plated on glass coverslips (Marienfeld #0115200) were washed once with PBS and fixed. For p-ERM and p-MLC2 staining, cells were fixed with 10% trichloroacetic acid (Sigma #T0699) for 10min at room temperature before extensive washings with TBS (20mM Tris-HCl pH 7.5, 154 mM NaCl, 2mM EGTA, 2mM MgCl_2_). For cytoskeleton staining, cells were fixed with 4% paraformaldehyde (Cederlane #043368-9M) for 30min. Cells were then permeabilized with 0,02% saponin (Amresco #0163) and blocked with 2% BSA (Bioshop #ALB001.250) for 1h. The antibodies used were the following: rabbit anti-phospho-ERM (1:500, overnight)^3^, mouse anti-phospho-MLC2 (Ser19) (1:50, overnight, Cell signaling #3675S), goat anti-rabbit Alexa Fluor 488-conjugated secondary antibody (1:200, 1h, Invitrogen #A11070), goat anti-mouse Alexa Fluor 488-conjugated secondary antibody (1:200, 1h, Invitrogen #A21235), phalloidin Texas red (1:100, 1h, Life technologies #T7471) and anti-alpha-Tubulin-FITC (1:200, 1h, Millipore Sigma #F2168). Coverslips were mounted in Vectashield medium with DAPI (Vector Laboratories #H-1200). Images were acquired using a LSM700 confocal microscope (Zeiss) with 63x objective. For cell sphericity measurements, orthogonal views were extracted using Zen software (Zeiss). Blue lines on pictures symbolize the adhesion surface (coverslip). P-ERM staining and cell sphericity were quantified using ImageJ software (NIH).

### In vitro kinase assay

In vitro kinase assays were performed as previously described with slight modifications^18^. Briefly, endogenous SLK from HEK293T cell lysate was immunoprecipitated using rabbit anti-SLK antibody (Bethyl #A300-499A) incubated for 1h at 4°C followed by incubation with protein A-sepharose beads (GE Healthcare #GE17-0780-01) for 2h at 4°C. Beads were then washed three times with TLB and three times with kinase reaction buffer (KRB) (50 mM Tris-HCl pH 7.5, 100 mM NaCl, 6 mM MgCl_2_ and 1 mM MnCl_2_). Beads were then resuspended in 15µl KRB buffer supplemented with 2 mM DTT, 50 µM ATP, 1 mM Na_3_VO_4_, 5 mM NaF, 2.5 mM sodium pyrophosphate and 100µl 10xPhosSTOP (Roche, #04906845001) with purified recombinant GST or GST-Ezrin^479-585^ from BL21 bacteria. Beads were finally incubated at 30°C for 30min, and proteins were denatured using sample buffer. For in vitro kinase assay using recombinant purified SLK, SLK kinase enzyme system was purchased from Promega (#V4242). In vitro kinase assays were performed in 384-well low volume plates (Greiner #784075) in 5µl reaction volume. Kinase reactions were executed for 2h at room temperature in reaction buffer A (40 mM Tris pH 7.5, 20 mM MgCl_2_ and 0,1 mg/ml BSA supplemented with 50 µM DTT) with 100 ng purified SLK, 0.5 µg histone H3 protein as substrate and 50 µM ATP. Kinase reactions were then stopped and processed using ADP-Glo kinase assay kit (Promega #V6930) as indicated in the manual. Briefly, 5 µl of ADP-Glo reagent was added for 40 min at room temperature to stop the reaction, followed by 10 µl of Kinase Detection Agent for 30 min. Luminescence signals was measured with a Synergy NEO Microplate Reader (Biotek).

## Data analysis

All quantifications were performed using image J software (NIH) and analyzed using GraphPad PRISM software (GraphPad Software, La Jolla, CA, USA). All data are represented by the mean +/-s.d. of multiple independent experiments. Microscopy images were prepared using Image J software (NIH) and Photoshop (Adobe).

## FIGURES & LEGENDS

**Figure S1.**
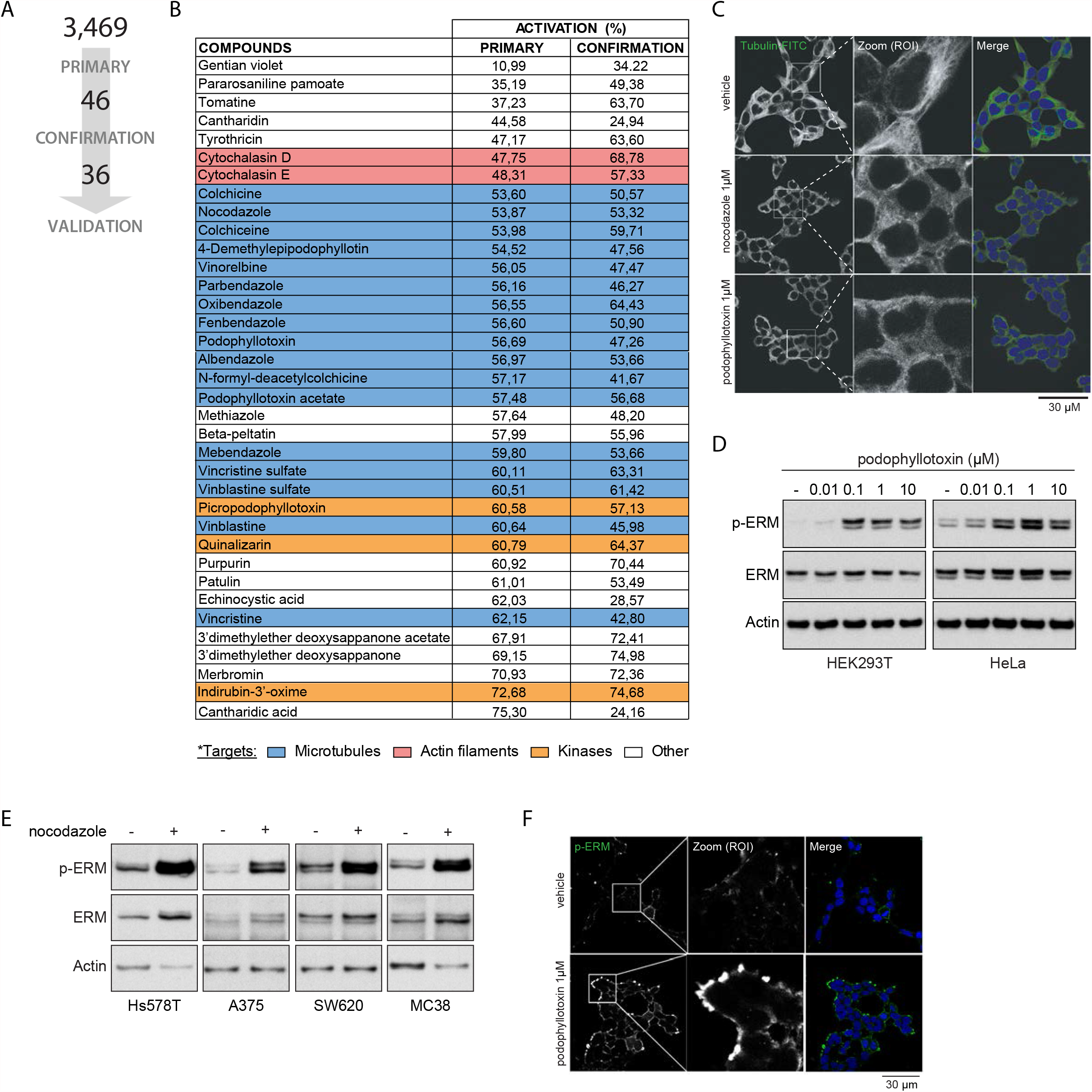
**(A)** Schematic representation of the number of hits identified after both the primary and the confirmation screens. **(B)** Table showing the targets of each compounds identified and validated in the chemical screen. BRET signals (in % compared to the vehicle) measured for each compound in the primary and confirmation screens are indicated in the columns 2 and 3. **(C)** Immunofluorescence of microtubule network of HEK293T cells treated with vehicle (DMSO), 1 µM nocodazole or 1 µM podophyllotoxin for 15 min. **(D)** Immunoblot of HEK293T (left) and HeLa (right) cells treated with increasing amount of podophyllotoxin for 15 min. **(E)** Immunoblot of Hs578T, A375, SX620 and MC38 cells treated with 1 µM nocodazole for 15 min. **(F)** Immunofluorescence of p-ERM staining in HEK293T cells treated with 1 µM podophyllotoxin for 15 min. Immunofluorescences (C, F) and immunoblots (D-E) are representative of three independent experiments.

**Figure S2.**
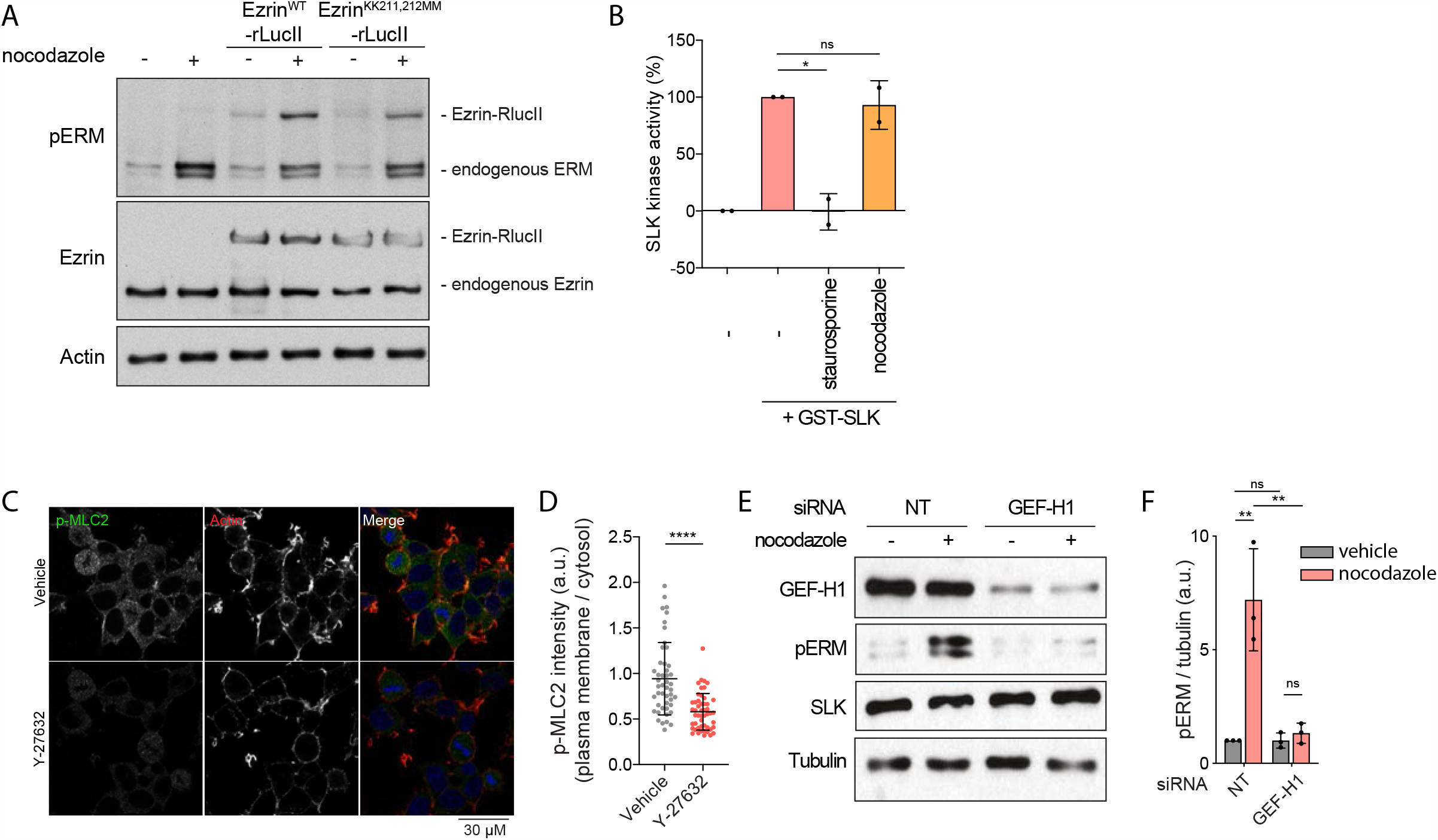
**(A)** Immunoblot of HEK293T cells transfected with Ezrin^WT^-rLucII or ^EzrinKK211,212MM^-rLucII and treated with vehicle (DMSO) or 1 µM nocodazole for 15 min. **(B)** SLK kinase activity measured by *in vitro* kinase activity in presence of purified GST-SLK pre-incubated with vehicle (DMSO), 100 nM staurosporine or 1 µM nocodazole for 15 min. **(C-D)** p-MLC2 (Ser19) and actin staining of HEK293T cells treated with vehicle or 10 µM Y-27632 for 4h (C). p-MLC2 signal intensity at the plasma membrane over cytosol is normalized to cells treated with vehicle (D). **(E-F)** Immunoblot of HEK293T cells depleted for GEF-H1 by siRNA and treated with either vehicle (DMSO) or 1 µM nocodazole for 15 min (A). P-ERM over tubulin signal was quantified and normalized to cells treated with non-target siRNA (NT) and treated with vehicle (B). Immunoblots (A, E) are representative of three independent experiments and immunofluorescence (C) is representative of two independent experiments. SLK kinase activity and p-ERM quantification represent the mean +/-s.d. of two or three independent experiments, respectively. Dots represent independent experiments (B, F) or independent cells (D). P values were calculated using Holm-Sidak’s multiple comparisons test with a single pooled variance (B, F) or unpaired t test (D). *, P < 0.05. **, P < 0.01. ****, P < 0,0001. ns, not significant.

**Figure S3.**
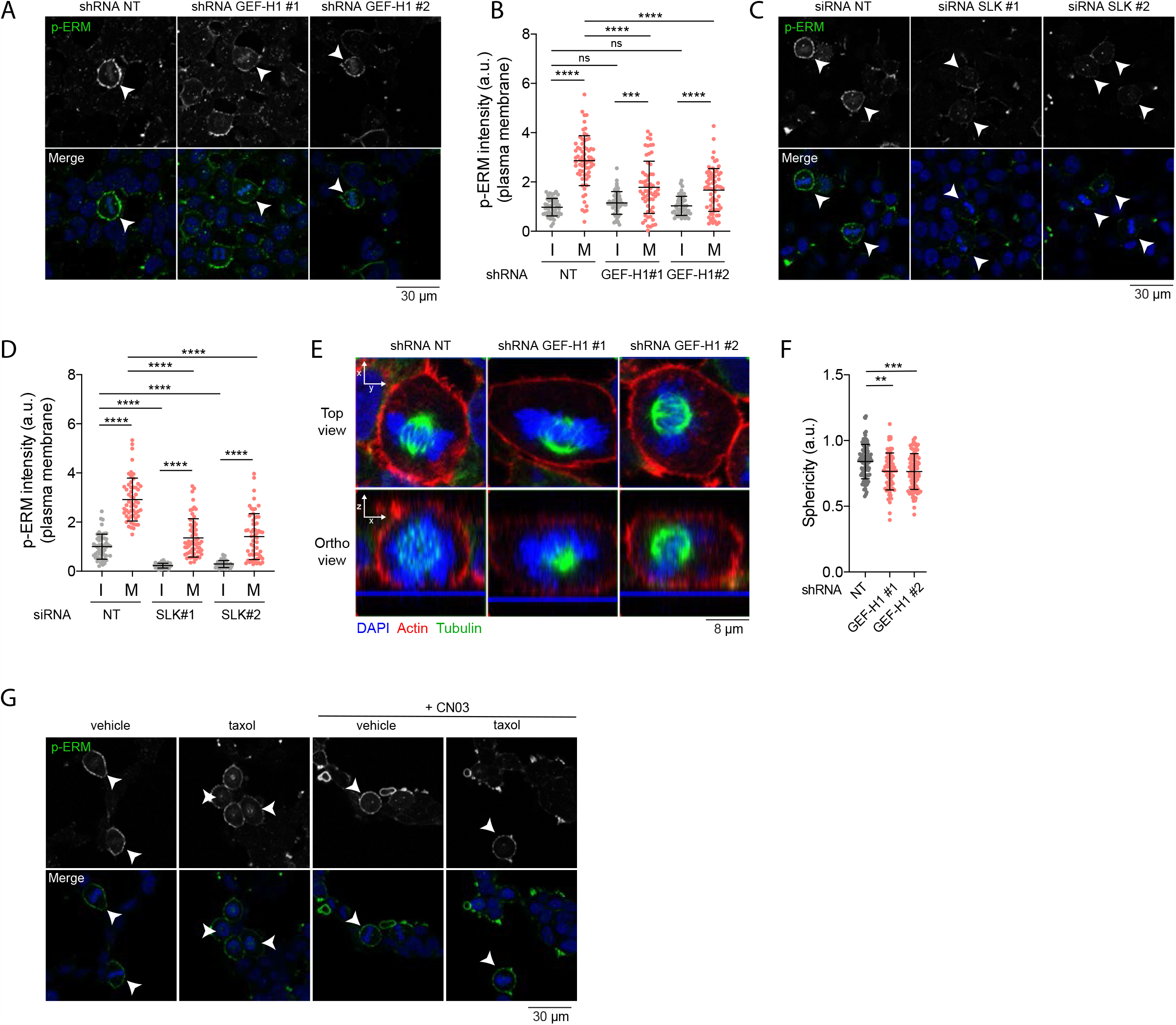
**(A-D)** p-ERM (white) and DAPI (blue) staining of HEK293T cells treated with non-target shRNA or 2 independent shRNA targeting GEF-H1 (A-B) or treated with non-target siRNA or 2 independent siRNA targeting SLK (C-D). Metaphase and interphase cells were identified based on DAPI staining (A, C) and p-ERM signal intensity at the plasma membrane is normalized to interphase cells treated with non-target shRNA/siRNA (NT). I = interphase, M = metaphase. **(E-F)** Sphericity of metaphase cells (F) was measured on immunofluorescence orthogonal reconstruction of Z stacks (E bottom) (F-actin, red; Tubulin, green; DAPI, blue) in HEK293T metaphase cells treated with non-target shRNA or 2 independent shRNA targeting GEF-H1. **(G)** p-ERM (white) and DAPI (blue) staining of HEK293T cells treated with vehicle (DMSO), 10nM taxol for 90 min and / or 1 µg/mL Rho activator II (CN03) for 1h. p-ERM signal intensity at the plasma membrane was quantified in Fig. 3Q. Immunofluorescences are representative of three independent experiments. P-ERM and sphericity quantifications represent the mean +/-s.d. of three independent experiments. Dots represent individual cells (B, D, F). P values were calculated using Sidak’s multiple comparisons test with a single pooled variance. **, P < 0.01. ***, P < 0.001. ****, P < 0.0001. ns, not significant.

**Figure S4.**
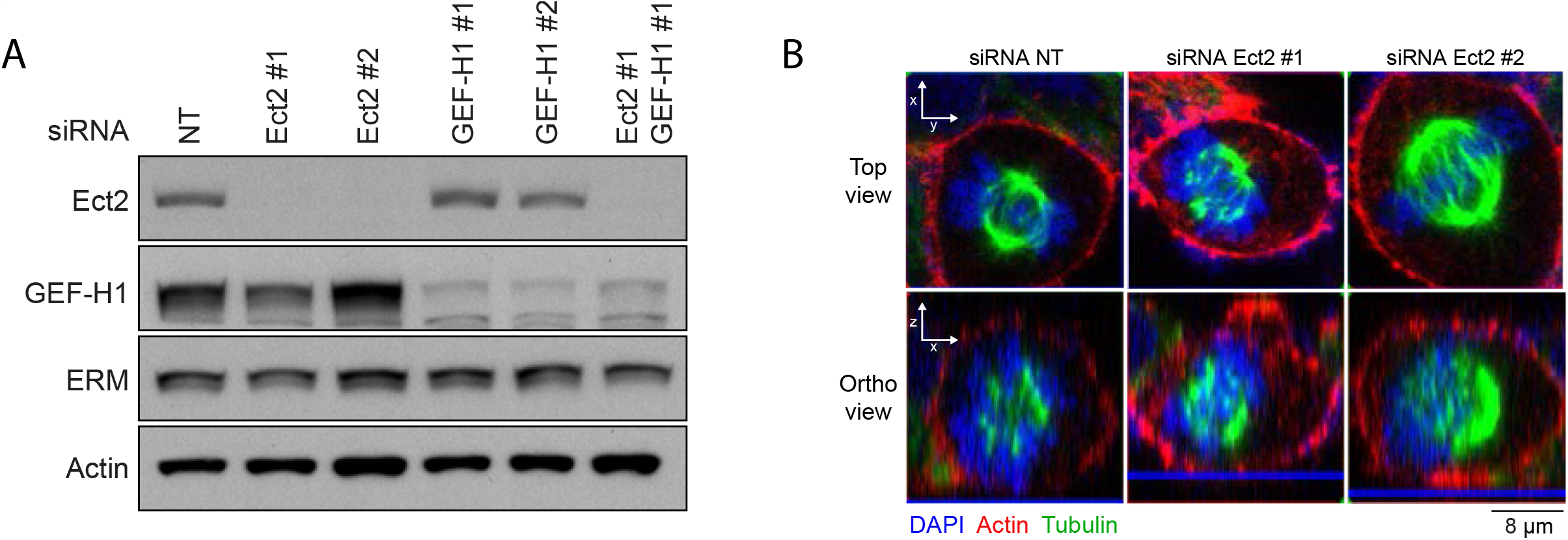
**(A)** Immunoblot of HEK293T cells treated with non-target siRNA (line 1), 2 independent siRNA targeting Ect2 (lines 2-3), 2 independent siRNA targeting GEF-H1 (lines 4-5) or with a siRNA targeting Ect2 plus a siRNA targeting GEF-H1 (line 6). **(B)** Immunofluorescence orthogonal reconstruction of Z stacks (bottom) (F-actin, red; Tubulin, green; DAPI, blue) in HEK293T metaphase cells treated with non-target siRNA or 2 independent siRNA targeting Ect2. Sphericity is measured in Fig 4C. Immunoblot (A) and immunofluorescence (B) are representative of three independent experiments.

